# Development of a vaccine against the newly emerging COVID-19 virus based on the receptor binding domain displayed on virus-like particles

**DOI:** 10.1101/2020.05.06.079830

**Authors:** Lisha Zha, Hongxin Zhao, Mona O. Mohsen, Liang Hong, Yuhang Zhou, Zehua Li, Hongquan Chen, Xuelan Liu, Xinyue Chang, Jie Zhang, Dong Li, Ke Wu, Monique Vogel, Martin F Bachmann, Junfeng Wang

## Abstract

The ongoing coronavirus COVID-19 pandemic is caused by a new coronavirus (SARS-CoV-2) with its origin in the city of Wuhan in China. From there it has been rapidly spreading to many cities inside and outside China. Nowadays more than 33 millions with deaths surpassing 1 million have been recorded worldwide thus representing a major health issue. Rapid development of a protective vaccine against COVID-19 is therefore of paramount importance. Here we demonstrated that recombinantly expressed receptor binding domain (RBD) of the spike protein homologous to SARS binds to ACE2, the viral receptor. Higly repetitive display of RBD on immunologically optimized virus-like particles derived from cucumber mosaic virus (CuMV_TT_) resulted in a vaccine candidate that induced high levels of specific antibodies in mice which were able to block binding of spike protein to ACE2 and potently neutralized COVID-19 virus *in vitro*.

## Introduction

COVID-19 is a disease caused by a novel coronavirus closely related to viruses causing SARS and MERS. As the disease caused by the other two viruses, COVID-19 mainly manifests symptoms in the lung and causes cough and fever (1). The disease COVID-19 is less severe then SARS and MERS, which is beneficial per se but leads to easier and more rapid spread of the virus, in particular due to infected individuals with very little sysmptoms (“spreaders”) and a potentially long incubation time (3 weeks) combined with viral shedding before disease onset (2). Indeed, COVID-19 has spread much further and more rapidly than the other two viruses. A vaccine with rapid onset of protection is therefore in high demand for the control of the pandemic that is currently taking its course. Different vaccine candidates are already in clinical trials all targeting either structural and enzymatic proteins or the spike proteins of SARS-CoV-2 (3). Most vaccine are either based on viral vectors (e.g. Adenovirus, AstraZeneca and J&J) or mRNA (Moderna, Curevac, Biontech), as such vaccine candidates can be rapidly produced under GMP conditions and no particular design is required. Both types of vaccines encoding the spike protein, the lipid nanoparticle encapsulated mRNA (4) and the recombinant adenovirus-based vaccines (5) have already been tested in individuals of different ages and are now in phase 2/3 clinical trials. Other vaccines are based on whole virus (live-attenuated or inactivated) and since mid-April 2020, a vaccine consisting of inactivated SARS-CoV-2 virus plus alum adjuvant is tested in China (6). However, it will be highly demanding to produce sufficient amounts of such a biosafety level-3 virus for the high global needs. Thus, an attractive possibility is to use inactivated nanoparticles or virus-like particles (VLPs) that have already delivered successful vaccines (7, 8). They can be engineered to display epitopes of foreign viruses on their surface, rendering those epitopes highly immunogenic. We have previously shown that antigens displayed on virus-like particles (VLP) induce high levels of antibodies in all species tested, including humans (9). More recently, we have developed an immunologically optimized VLP platform based on cumcumber mosaic virus. These CuMV_TT_ VLPs incorporate a universal T cell epitope derived from tetanus toxin providing pre-existing T cell help. In addition, these VLPs package bacterial RNA which is a ligand for toll-like receptor 7/8 and serves as potent adjuvants (10). Using antigens displayed on these VLPs, it was possible to induce high levels of specific antibodies in mice, rats, cats, dogs and horses and treat diseases such as atopic dermatitis in dogs or insect bite hypersensitivity in horses (10–12).

Here, we used CuMV_TT_-VLP as vaccine platform to generate a vaccine against the COVID-19 spike protein. The spike protein of COVID-19 is highly homologous to the spike protein of SARS and both viruses share the same receptor, which is angiotensin converting enzyme 2 (ACE2) (13, 14). The receptor binding domain (RBD) of the SARS spike protein binds to ACE2 and is an important target for neutralizing antibodies (15, 16). By analogy the RBD of COVID-19 spike protein may be expected to similarly be the target of neutralizing antibodies, blocking the interaction of the virus with its receptor. To this end we generated RBD-CuMV_TT_ vaccine by chemically coupling RBD of SARS-CoV-2 on the CuMV_TT_-VLP. We showed that the vaccine was highly immunogenic in mice and was able to generate antigen-specific antibodies with neutralizing effect on SARS-CoV-2.

## Material and Methods

### Protein expression and purification

The COVID-19 receptor-binding domain (RBD) and the N-terminal peptidase domain of human ACE2 were expressed using HEK293F cells (Invitrogen). The COVID-19 RBD (residues Arg319-Phe541) with an N-terminal IL-2 signal peptide for secretion and a Fc part (hinge-CH2-CH3) of human IgG1 for purification was inserted into pFUSE-mIgG1-Fc2 vector (Invitrogen, Thermo Fisher Scientific, MA, USA). The construct (RBD-huFc) was transformed into bacterial DH5α competent cells, and the extracted plasmid was then transfected into HEK293F cells at a density of 3×10^6^ cells/ml using PEI (Invitrogen, Thermo Fisher Scientific, MA, USA). The supernatant of cell culture containing the secreted RBD was harvested 72 h after infection, concentrated and buffer-exchanged to HBS (10 mM HEPES, pH 7.2, 150 mM NaCl). RBD was captured by protein A resin (GE Healthcare) and eluted with Gly-HCl buffer pH 2.2. Fractions containing RBD were collected and neutralized to pH 7.0 with 1M Tris. For ELISA coating, RBD was cleaved from the Fc part using thrombin(Beijing Solarbio Science & Technology Co., Ltd.) as described in the manufacturer’s manual.

The human ACE2 (residues Ser19-Ser741) with an N-terminal IL-2 signal peptide for secretion and a C-terminal 6×His tag for purification was inserted into pFUSE-vector (Invitrogen, Thermo Fischer Scientific, MA, USA). The human ACE2 was expressed by essentially the same protocol used for the COVID-19 RBD. ACE2 was captured by Ni-NTA resin (GE Healthcare) and eluted with 500 mM imidazole in HBS buffer. ACE2 was then purified by gel filtration chromatography using the Superdex 200 column (GE Healthcare) pre-equilibrated with HBS buffer. Fractions containing ACE2 were collected.

### Serum Competitive ELISA

The antibody competitive binding activities of the serum were assayed by ELISA. ACE2 (1 μg/ml) was incubated in 96-well plate overnight at 4 °C. After incubation, the plate was blocked with 2% BSA for 2h at 37°C and then was washed five times with PBS containing 0.05% Tween 20. BSA was used as negative control followed by the addition of a mixture of 40-fold diluted serum and RBD-huFc (0.15 μg/ml) followed by incubation for 30 min with gentle shaking at 37°C. Plates were washed five times with PBS containing 0.05% Tween 20 (PBST) followed by 100 μl of horseradish peroxidase/anti-mFc antibody conjugate (diluted 1:5000 in PBT buffer), incubated 30 min with gentle shaking. Plates were washed five times PBT buffer and developed with 100 μl of freshly prepared 3,3’,5,5’-Tetramethylbenzidine (TMB) substrate. Reaction was stopped with 100 μl of 1.0 M H3PO4 and read spectrophotometrically at 450 nm in a microtiter plate reader.

### Production of CuMV_TT_-VLP

The production of CuMV_TT_-VLP was described in detail in Zeltins et al.^9^. Briefly, *E coli* C2566 cells (New England Biolabs, Ipswich, MA) were transformed with the CuMV_TT_ coat protein (CP) gene-containing plasmid pET CuMV_TT_. The expression was induced with 0.2 mM isopropyl-β-D-thiogalactopyranoside (IPTG). The resulting biomass was collected by low-speed centrifugation and was frozen at −20°C. After thawing on ice, the cells were suspended in the buffer containing 50 mM sodium citrate, 5 mM sodium borate, 5 mM EDTA, and 5 mM 2-Mercaptoethanol (pH 9.0, buffer A) and were disrupted by ultrasonic treatment. Insoluble proteins and cell debris were removed by centrifugation (13,000 rpm, 30 minutes at 5°C, JA-30.50Ti rotor, Beckman, Palo Alto, CA). The soluble CuMV_TT_ CP protein in clarified lysate was pelleted using saturated ammonium sulfate (1:1, vol/vol) overnight at 4°C. Soluble CuMV-CP-containing protein solution was separated from the cellular proteins by ultracentrifugation in a sucrose gradient (20%-60% sucrose; ultracentrifugation at 25,000 rpm for 6 hours at 5°C [SW28 rotor, Beckman]). After dialysis CuMV_TT_-containing gradient fractions, VLPs were concentrated using ultracentrifuge (TLA100.3 rotor, Beckman; at 72,000 rpm for 1 hour, +5°C) or by ultrafiltration using Amicon Ultra 15 (100 kDa; Merck-Millipore, Cork, Ireland).

### Generation of the vaccine CuMV_TT_-RBD

The RBD was conjugated to CuMV_TT_ using the cross-linker Succinimidyl 6-(beta-maleimidopropionamido) hexanoate (SMPH) (Thermo Fisher Scientific, MA) at 10-molar excess for 60 minutes at 23°C. The coupling reactions were performed with 0.3x molar excess of RBD, 0.3x RBD, or equal molar amount of RBD regarding the CuMV_TTS_ by shaking at 23°C for 3 hours at 1200 rpm on DSG Titertek (Flow Laboratories, Irvine, UK). Unreacted SMPH and RBD proteins were removed using Amicon-Ultra 0.5, 100K (Merck-Millipore, Burlington, Ma). VLP samples were centrifuged for 2 minutes at 14,000 rpm for measurement on ND-1000. Coupling efficiency was calculated by densitometry (as previously described for IL17A-CuMV_TT_ vaccine^9^), with a result of approximately 20% to 30% efficiency.

### Mice

BALB/c mice at the age of 6 weeks were purchased from Beijing Vital River Laboratory Animal Technology Co., Ltd and kept at the animal facility of Anhui Agricultural University. All animal experiments were performed in accordance with the National Animal Protection Guidelines and approved by the Chinese Association for Laboratory Animal Sciences.

### Vaccination

Six-week-old naive BALB/c mice (3 mice per group) were immunized by subcutaneous injection with 50 μg of either RBD-CuMV_TT_, RBD or CuMV_TT_ as control. The boosts of the vaccination were conducted at 2^nd^ and 3^rd^ week. Serum was collected for ELISA analysis 1 week after every vaccination.

### Generation of pseudovirus

Pseudovirus expressing the SARS-CoV-2/COVID-19 spike protein were produced by lentivirus second-generation packing system^15^. Plasmids of pwpxl-luc, HIV-1 PSD and pCMV3 containing the SARS-CoV-2/COVID-19 Spike gene were co-transfected into 7 x 10^5^ 293F cells using Sinofection (Sino Biological, Beijing, China). The medium was replaced with fresh DMEM containing 10% FBS after overnight incubation. Supernatants containing pseudovirus were collected 48 h and 72h after transfection and filtered using a 0.45-μm filter syringe. All filtered supernatants were collected together and stored at −80 °C until use.

### Titration of pseudovirus

The 293T-ACE2 cells which stably express ACE2 receptors on the cell membrane were prepared by transfection of ACE2 gene into 293T cells using lentivirus system. Pseudoviruses prepared above were added to the 293T-ACE2 cells (3 × 10^4^ cells/well) with 100 μl polybrene (16 μg/ml). After 48 h, the infection was monitored using the Luciferase Assay System (Promega, Madison, USA). Titer was calculated based on serial dilutions of pseudovirus.

### Neutralization Assay

The mouse serum samples (2 μl) were diluted to 1:10, 1:40, 1:160, 1:640 and 1:2560 respectively, and then mixed with an equal volume of pseudovirus stock. After incubation at 37°C for 1 h, the mixture was inoculated on the 293T-ACE2 cells (3 x 10^4^ cells/well). At the same time, pseudovirus+DMEM medium was set as a positive control and DMEM medium only was set as a negative control. After the cells were incubated for 72 hours, serum neutralization was measured by luciferase activity of infected pseudovirus. A cut-off of >80% was used as to detetermine neutralizing titer.

## Results

### Generation of the RBD-CuMV_TT_ vaccine

To generate a COVID-19 vaccine candidate, we displayed the RBD domain of SARS-CoV-2 spike protein on CuMV_TT_ (Fig. 1a). To this end, we gene-synthesized the COVID-19 RBD domain and fused it to an Fc of human IgG1 for better expression. As shown by sandwich ELISA, the protein bound efficiently to the viral receptor ACE2 (Fig. 1b). In a next step, the protein was chemically coupled to the surface of CuMV_TT_ using the well established chemical cross-linkers SATA and SMPH (10). SDS-Page and Western Blotting(HRP-GOAT-anti-Mouse IgG, Shanghai Huaxin Biotechnology Co.,Ltd.) confirmed efficient coupling of the RBD-fusion molecule to the CuMV_TT_, resulting in the vaccine candidate RBD-CuMV_TT_ (Fig. 1c, d).

**Fig. 1.**
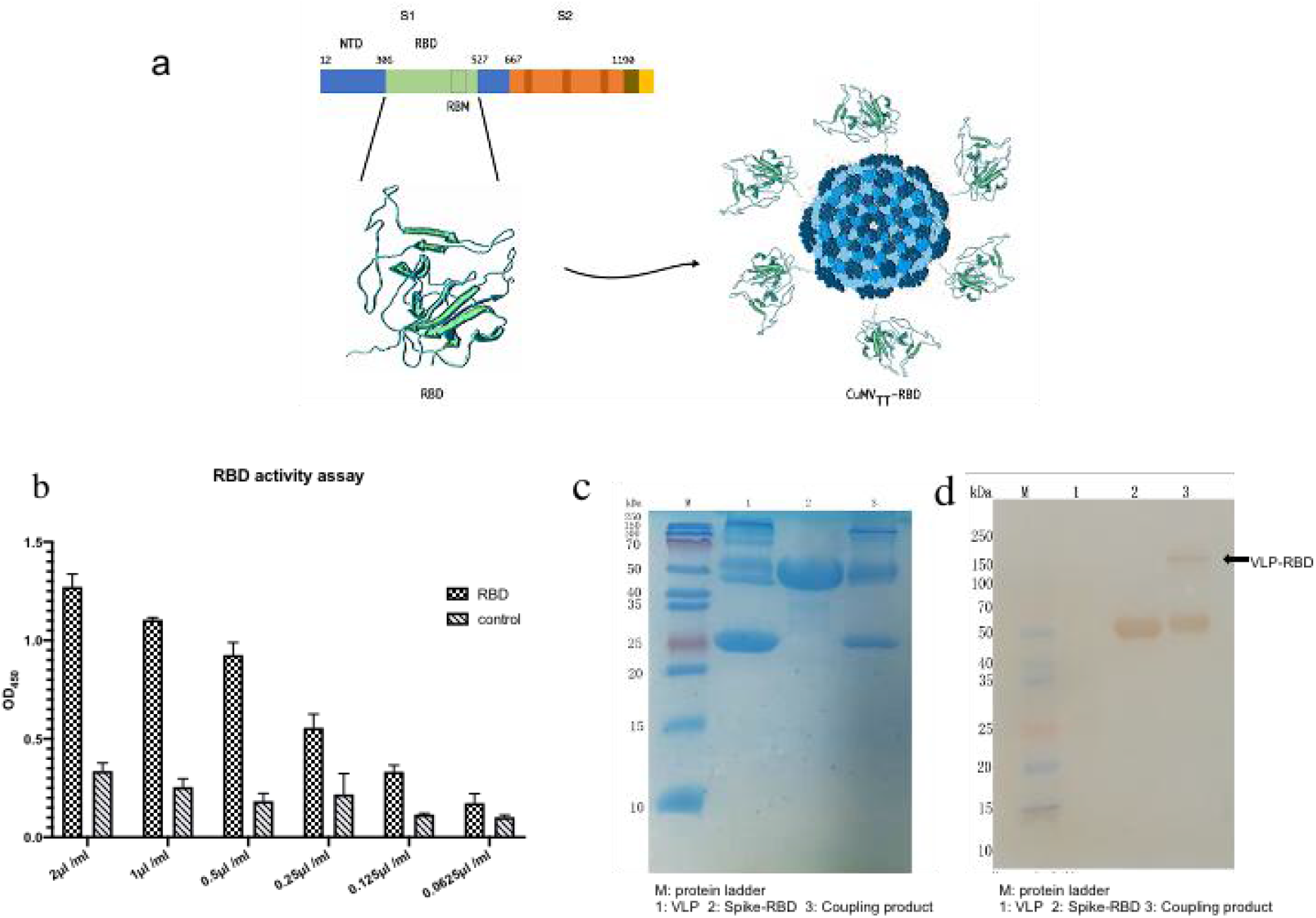
Coupling of Spike-RBD on CuMV_TT_-VLP. A) Outline of vaccine design strategy. B) Recombinantly expression and binding of RBD to ACE2 was demonstrated by ELISA. Different concentrations of RBD were tested on immobilized ACE2. C,D) RBD was displayed on the surface of VLP via SATA and SMPH chemical cross-linker, resulting in a coupling band (150kd) as shown by C) SDS-PAGE and D) Western blotting.

### Immunogenicity of RBD-CuMV_TT_ vaccine

To test the immunogenicity of the vaccine candidate, mice were immunized three times (weekly schedule) with the RBD-huFc fusion molecule alone or conjugated to the surface of CuMV_TT_ formulated in Montanide adjuvants. Serum samples were collected at weeks 1, 2 and the detection of anti-RBD antibodies were determined by ELISA using RBD as coating antigen. Our data demonstrated that coupling RBD to VLPs dramatically increased the immunogenicity of the RBD. As shown in Fig. 2a-c antibody titers are substantially increased in mice immunized with RBD-CuMV_TT_ after each vaccination. Immunization of mice with RBD alone only induces only weak antibody response that did not increase after each vaccination.

**Fig. 2.**
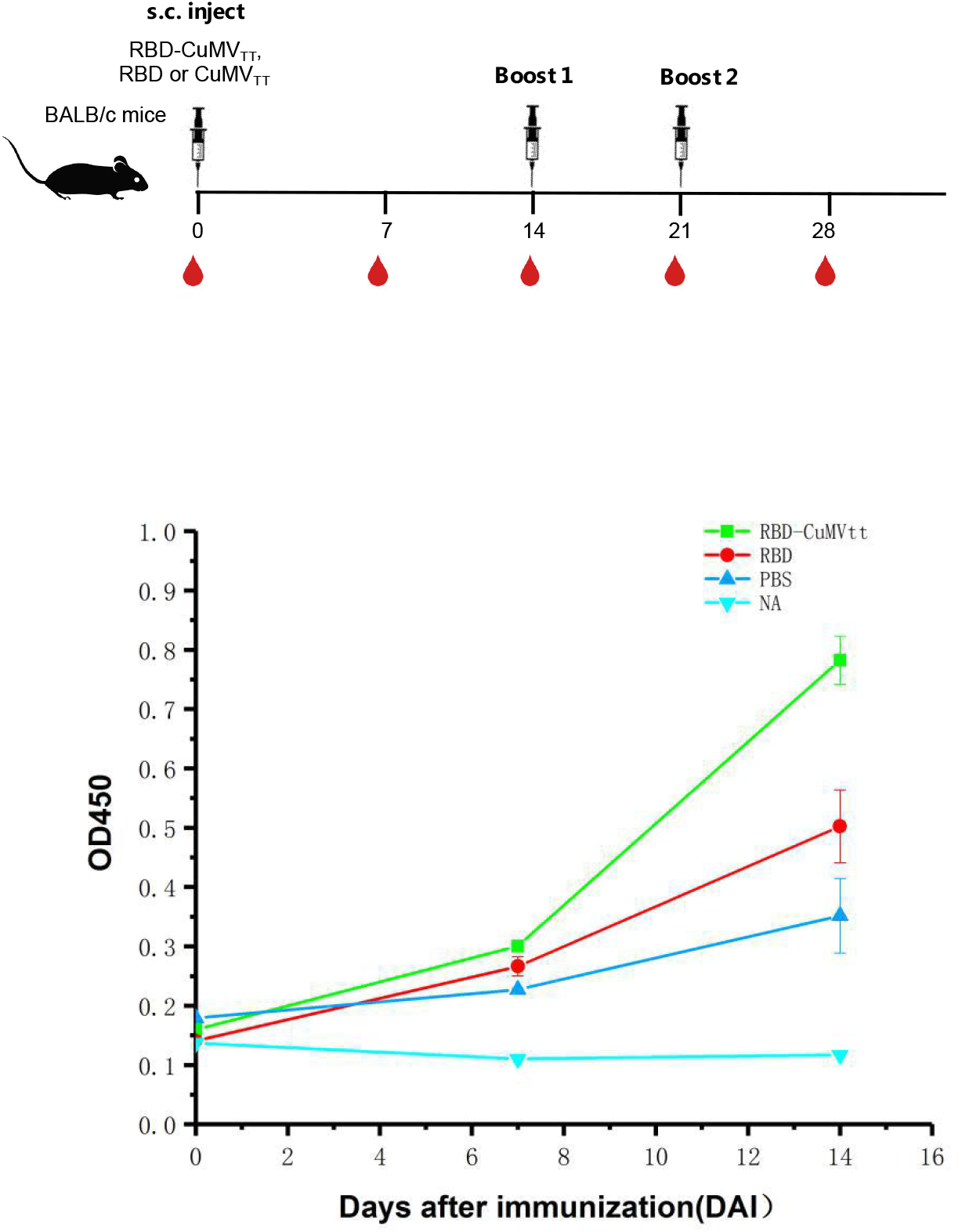
RBD-CuMV_TT_ induces high specific antibody titers. Mice (3 per group) were immunized with 10 μg of RBD-CuMV_TT_, RBD alone or CuMV_TT_ alone formulated in Montanide on days 0, 7 and 14. Serum was harvested on day 7 (A) and14 (B) and tested by ELISA on recombinant RBD. Note that strong antibody responses could be observed 14 day after immunization with RBD-CuMV_TT_.

Next, to determine the epitopes recognized by the anti-RBD antibodies we measured the competition of the antibodies with ACE2 for the binding to RBD by ELISA. HuACE2 was immobilised on the plates followed by the addition of the tested antibodies in the presence of RBD-huFc. As shown in Fig. 3, sera of mice immunized with RBD-CuMVTT obtained after two boosts (day 21) were able to strongly inhibit RBD binding to ACE2. For comparison, sera from mice immunized with RBD alone or with CuMVTT did not show any significant competition for the binding to RBD. These results indicate that immunization with RBD-CuMVTT induce anti-RBD antibodies recognizing an epitope that overlaps with the ACE2 binding of COVID-19 RBD.

**Fig. 3.**
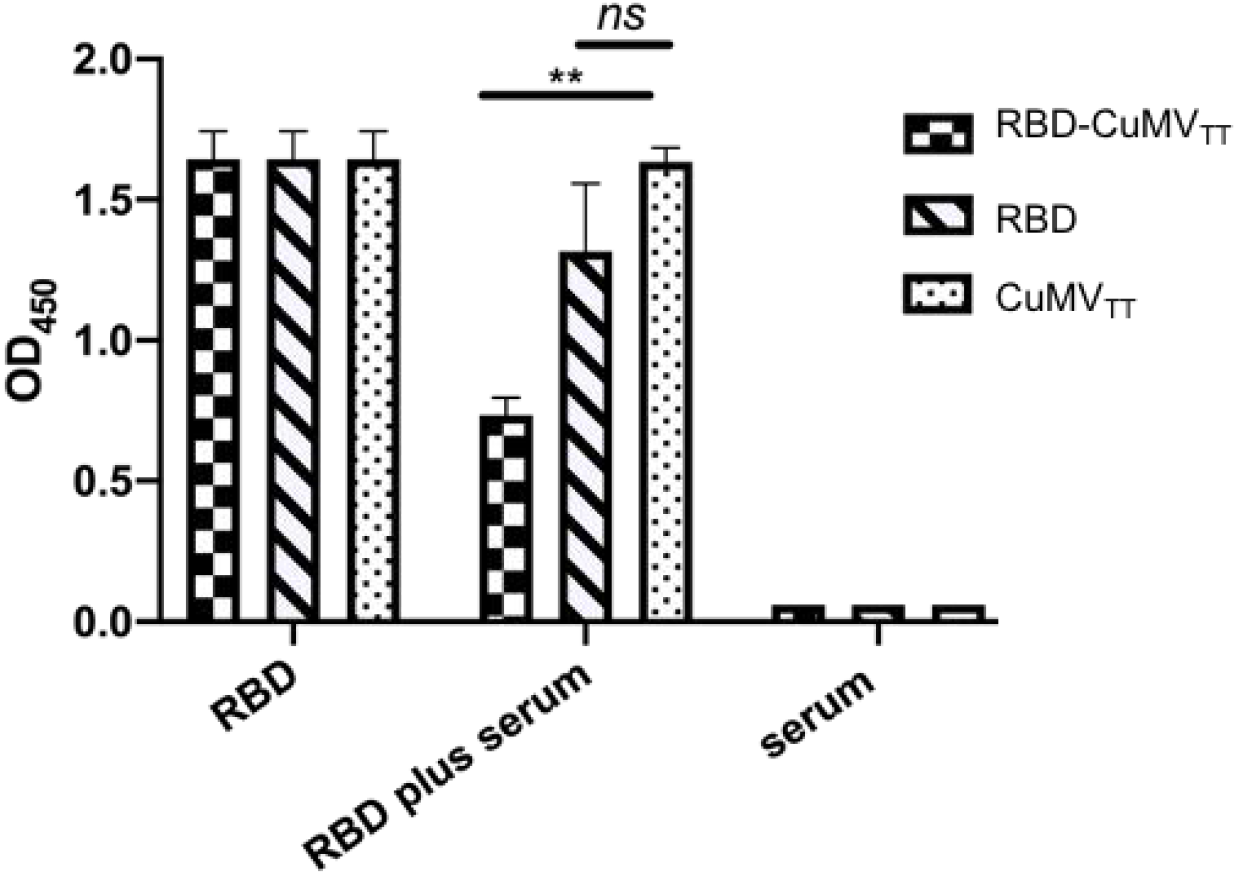
Immunization with RBD-CuMV_TT_-induces antibodies that block binding of RBD to ACE2. Mouse sera (1:40 dilution, d21) after 3rd vaccination were incubated with RBD at a concentration of 0.15 g/ml for 1 hour then incubated on the ACE2 coated plate. RBD binding was determined by ELISA.

### Neutralizing activity of the anti-RBD antibodies

As the infection of the cells by SARS-CoV-2 is mediated through the binding of the spike protein to the ACE2 the best correlation of protection is viral neutralization. To this end we generated pseudotyped retroviruses^12^ expressing the COVID-19 spike protein and luciferase for quantification of infection (Fig. 4a). Using these viruses, the neutralizing capacity of the sera from immunized mice was assessed on ACE2-transfected cells (Fig. 4b), directly demonstrating high anti-viral neutralizing activity of the induced antibodies. Hence, the RBD-CuMV_TT_ vaccine candidate was able to induce high levels of COVID-19 virus neutralizing antibodies whereas sera from mice immunized with RBD alone or with CuMVTT did not show any neutralizing activity. From these experiments we conclude that the CuMV_TT_ based vaccine based on highly efficient expression systems and chemical conjugation technologies, is able to induce high titers of neutralizing antibodies that can interfere with the binding of the virus to the human cells.

**Fig. 4.**
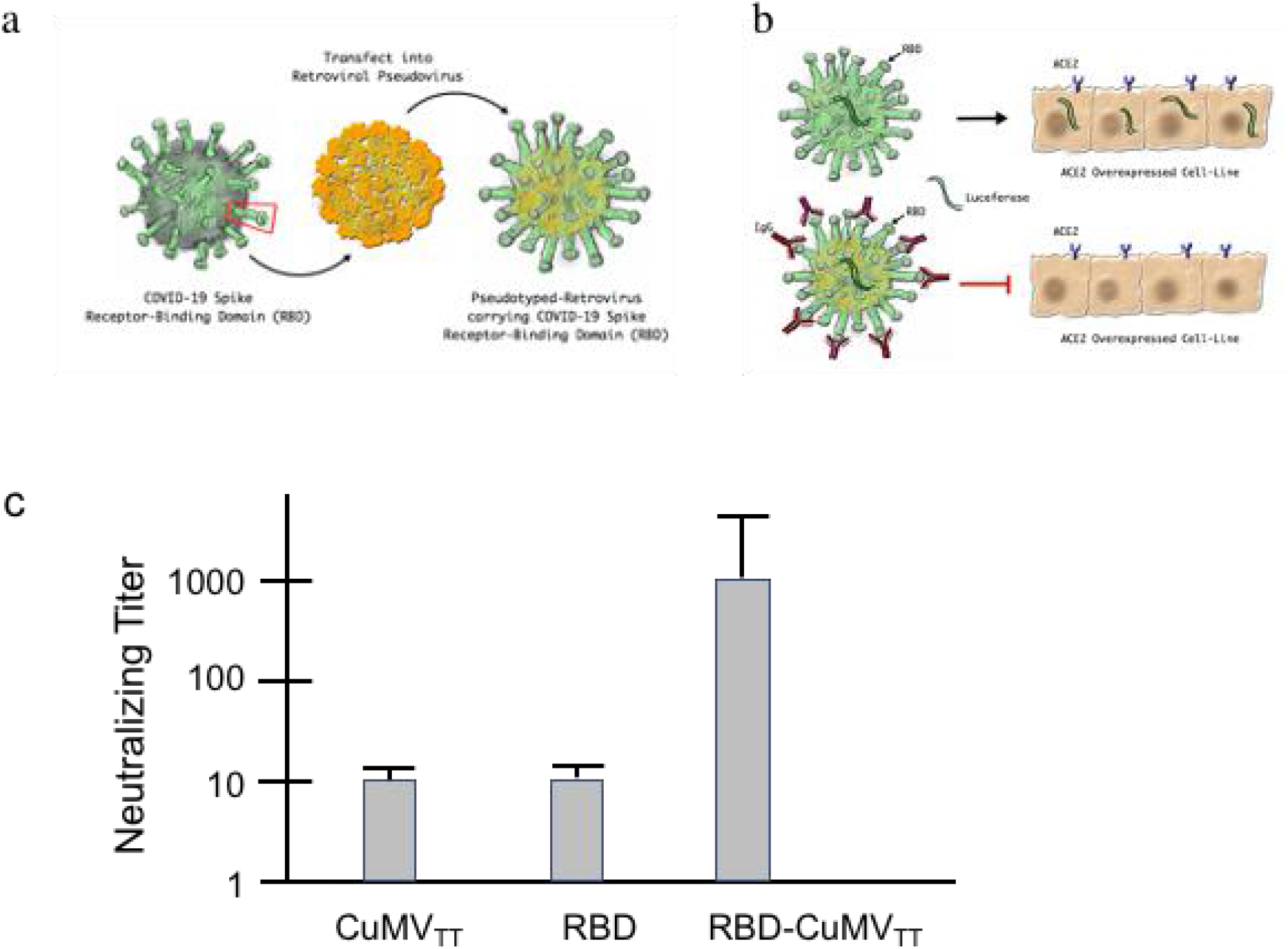
Serum neutralization assay: Pseudovirus expressing the COVID-19 spike protein was generated by co-transfection of the plasmids of luciferase-expressing HIV pseudoparticles into HEK 293 T cells using second-generation lentiviral packaging system. A,B) Schematic presentation of the assay principle. C) Pseudovirus expressing the SARS-CoV-2 spike protein can infect the cells by ACE2 receptor and antibodies induced by RBD-CuMV_TT_ (day 21) potently neutralize the virus.

## Discussion

The immune response to the SARS-CoV-2 infection is initiated by innate immune activation followed by antigen-specific T and B cell responses (17). The most important mechanism protecting against reinfection is presence of virus neutralizing antibodies which is similar for almost all viruses causing acute disease followed by pathogen clearance. Here we demonstrated that display of RBD-domain in CuMV_TT_ VLPs induces high levels of IgG antibodies that bind recombinant RBD in ELISA experiments. These antibodies were functionally validated by demonstrating that induced antibodies can block binding of RBD to ACE2, the receptor for SARS-CoV-2. Such competitive inhibition is an important feature for viral neutralization (13, 14, 18). Indeed, the induced antibodies were able to potently neutralized lentiviruses pseudotyped with the spike of SARS-CoV-2.

Antibody, particular neutralizing antibody responses to Coronaviruses are often weak and short lived, in particular in patients with little or no symptoms (19). Although many patients generate overall SARS-CoV-2 specific antibody-responses, neutralizing antibodies often remain at low titers and appear late. Of greatest concern may be that some patients may not generate long-lasting antibodies at all (20). There is accumulating evidence that protection from reinfection can be very short lived as demonstrated by some patients which experienced COVID-19 twice, despite a proven virus free interval in between (21). Thus, on average, SARS-CoV-2 triggers a curiously weak and short-lived response while most other acute disease-causing viruses induce long-lived neutralizing antibody responses (22). Similarly, most attenuated, replication competent viral vaccines induce long-lived antibody responses after a single shot in the absence of disease.

The surface geometry of SARS-CoV-2 may offer an explanation for the unexpected shortlived antibody responses. Most viruses exhibit surface epitopes in a rigid manner, distanced by about 5 nm (8). This is optimal for induction of potent and long-lived immune responses and known as a pathogen-associated structural pattern (23).

Indeed, such viral particles efficiently cross-link B cell receptors (24, 25) and are recognized by natural IgM, causing activation of the classical pathway of complement. This causes binding of viral particles to complement receptors CD21 followed by B cell-dependent migration to and deposition on follicular dendritic cells resulting in efficient germinal center formation (26). Importantly, engagement of CD21 by antigen bound to complement-fragments allows induction of long-lived plasma cells, resulting in durable antibody responses (27). In contrast to these average viruses, SARS-CoV-2 is larger and the RBD epitopes are spaced by much longer distances in the range of 25 nm. Hence, SARS-CoV-2 may avoid strong and long-lasting antibody responses by diluting out the neutralizing epitopes.

The here presented strategy may allow to overcome this problem. Indeed, by displaying the RBD domain in a repetitive fashion on immunologically optimized CuMV_TT_ VLPs, one is able to shorten the distances to the optimal 5 nm, resulting in strong antibody responses. Thus, putting RBD in a more immunogenic context may allow to enhance RBD-specific neutralizing antibody responses as desired for an efficient vaccine.

